# Conservation vs. Wild-Animal Suffering : how can population dynamics help?

**DOI:** 10.1101/2022.10.10.511528

**Authors:** Nicolas Salliou, Paula Mayer, Alexandre Baron

## Abstract

Conservation and ethical consideration for animal welfare in the wild appear to be synergetic because they both care for non-human animals. However, many practices such as culling seem to achieve conservation purposes but at the cost of producing a lot of wild-animal suffering, antagonizing conservationists and animal rights advocates. To explore this tension, we model the suffering of animals in wild ecosystems by resorting to classical population dynamics equations and using death rates as a metric of suffering. Our results show that, depending on the structure and parameters of the ecosystem, animal rights advocates and conservationists can have either opposing or compatible interests, where conserving species can go hand in hand with reducing the overall suffering. These models contribute to the concrete question of how to cope with suffering in the wild and may help ecosystem managers who are regularly confronted with interventions in the wild.

## 1 Introduction

### Non human animal sentience

Animal advocacy has a long-standing history and dates as far back as ancient Greece in Europe. Pythagoras believed that killing and eating non-human animals spoiled the soul. However, Humanism, as well as most religions have focused on the specificity – and ultimately the superiority – of the human being over non-human animals (De Fontenay, 2014; Larue, 2015). This has translated into the way we have considered and treated animals for centuries, that is, as beings devoid of complex sentience. For several decades now, we have gradually garnered a considerable amount of knowledge about non-human animal sentience and the frontier between humans and non-humans has slowly started to fade. It is perhaps most apparent for one of the key attributes traditionally believed to be unique to humans: consciousness. In particular, the classical mirror self-recognition test, considered to be a founding trial of self-awareness, has been successfully passed by many animals including elephants, apes, magpies, and dolphins (Proctor, 2012). Consciousness suggests that the experience of positive and negative emotions by these animals is likely to be as meaningful as it is for humans. In July 2012, the Cambridge convention on consciousness that gathered cognitive neuroscientists, neuropharmacologists, neurophysiologists, neuroanatomists and scientists from several other fields from around the world, declared that a large variety of non-human animals, including all mammals, birds, and many other creatures, such as octopuses, possess the neurological substrate for consciousness and thus sentience (Low et al., 2012).

### Sentience, suffering and ethical concern for animals

In 1973, Peter Singer defended the idea that the capacity for sentience is sufficient to consider animal suffering from an ethical perspective in the framework of utilitarianism (Singer, 1973). The equal consideration for human and non-human suffering, already defended by the father of utilitarianism, Jeremy Bentham (Kniess, 2019), paved the way for a reconsideration of the way we treat domesticated animals, that is, by inflicting a great deal of suffering to benefit from their flesh, skin, milk, etc.

Even if we were to hypothesize that there is a weak probability that non-human animals have an equal capacity to feel pain, the sheer scale of potential suffering – hundreds of billions of animals are killed each year – makes a strong ethical case for the consideration of animal suffering, or else there is a high probability that it would be ethically disastrous (Huemer, 2019). Increasingly, modern moral philosophy has started to consider animals as selves and some contemporary political philosophers make a strong point in saying that they should be included in the community of persons and that as a result, they should be given universal basic rights (Regan, 2004; Donaldson and Kymlicka, 2011). This recognition of animals as persons mostly focuses on domesticated animals because humans have obviously a direct role in their lives. Recently, there has been increasing consideration for animals living in the wild.

### Recognizing wild-animal suffering (WAS)

While domesticated animal suffering and Animal Rights (AR) theories have gained considerable attention in recent years, the case for wild-animal suffering (WAS) has remained more confidential, but is nonetheless generating tension between the ecological and ethical perspectives on wildlife.

From an ecological standpoint, key studies have illustrated the role that the ecology of fear plays in well-functioning ecosystems. For instance, they have shown the important role that wolves play on the overall biodiversity of the Yellowstone National Park (Ripple and Beschta, 2004). Similarly, a lack of large predators such as grizzlies or wolves has been correlated to higher moose density and lower levels of avian biodiversity (Berger et al., 2001). The Yellowstone National Park has become a symbol of the positive effects of predation from an ecological perspective. Similarly, human predation is said to be justified for regulation purposes. It can maintain healthy populations by avoiding overpopulation, hunger, disease and habitat overexploitation. However, Zanette *et al*. (Zanette et al., 2019) have recently shown that the fear of predators generates changes in the brains and behavior of preys that resemble post-traumatic stress disorder (PTSD), which results in a reduction in the fecundity and survival of these animals. This shows that even though the ecology of fear can be beneficial from an ecological perspective, it might actually degrade the welfare of animal individuals.

On the Animal Rights (AR) side, traditional theories have promoted a *hands-off* approach of sorts. In other words, humans should stop harming wild animals to start with, and then leave them alone. In 1983, Tom Regan defended the idea that non-human animals are entitled to certain basic rights, including the right to live their own life and as a result humans should “let wildlife be” (Regan, 2004). Similarly, Donaldson and Kymlicka brought forward the idea that wild animals are sovereign in their territories and as a consequence, humans should limit their interactions as we would do for a foreign nation by respecting its own laws (Donaldson and Kymlicka, 2011). Canadian road signs stating “You’re in Bear country” follow some notion of wild-animal sovereignty. However, the foundations of wild-animal sovereignty have been questioned recently due to the lack of autonomy and morality of non-human animals (Ladwig, 2015), which would constitute prerequisites to sovereignty. Some authors question the sovereignty model because many wild animal communities are high growth rate (*r*) species (r-selected species) that produce lots of expendable offsprings, resulting in numerous deaths. This puts these communities in a state of constant crisis and as a result, they may be viewed as failed states rendering their sovereignty illegitimate (Horta, 2010, 2013). Finally, Palmer defended the idea that WAS should be considered in proportion to the involvement humans have in the suffering of wild animals as would be the case for road kills, or human generated forest fires (Palmer, 2015).

### Intervention to deal with WAS

In the previous section we showed that WAS is an issue that needs to be acknowledged. In this section, we show the current literature about intervention to limit WAS and introduce the key conditions before any action in real ecosystem. Aiding a wild animal in distress is regarded as particularly humane and compassionate. There are many accounts of such stories published on online plateforms, from the rescuing of koalas trapped in Australian bushfires to saving deers from drowning in lakes. These stories usually generate positive reactions for their display of empathy and kindness. While such isolated actions seem to remain unquestioned, probably due to their small scale, a mass intervention in wilderness to reduce WAS more systematically is usually met with skepticism (Coghlan and Cardilini, 2022).

However, in practice, fairly wide scale interventions in places usually considered wild, like National Parks, are extremely common. They may take the form of assisted reproduction of endangered species, armed squads against poachers, feeding (Dubois and Fraser, 2013), mass vaccination, culling to regulate species (Wilson et al., 2015), etc. For example, there have been vast vaccination campaigns of foxes against rabies, but this only collaterally reduced animal suffering as human health was the primary concern (Sidwa et al., 2005). It shows that large scale actions are possible and can achieve a reduction of WAS. While intervention in wild populations is extremely common and is usually socially well accepted, the idea of intervening to reduce WAS usually garners opposition and, to our knowledge, there are currently no large scale initiatives that primarily aim at reducing WAS.

Scholars in favor of intervention against WAS can be quite radical in their proposals. Some authors have defended the idea of using gene editing technology to change the biology of r-selected species and move them towards K-selected strategies, that is strategies where the species tend to live at high densities close to the carrying capacity (*κ*) of the environment (Johannsen, 2017). In contrast to r-selected species, that usually have fewer and well-cared after offspring. Other authors explore distant futures when predation could be reprogrammed through education or gene editing towards non-predatory behaviors (Pearce, 2009), or carnivorous species would be given in-vitro meat, following a form of “high-tech Jainism” approach (Pearce, 2016). Another idea put forward is selective extinction, where predators would die out using contraceptive strategies (Pearce, 2009).

Before any decision is made, many scholars have made clear that a pre-condition for any intervention to reduce WAS, is to have the capacity to predict the effects of intervention by using models (Delon and Purves, 2018). Some authors argue that intervention to reduce WAS is legitimate under the condition that it is feasible and that a positive outcome can be predicted (Faria et al., 2016; Torres, 2015). Another perspective aims at maintaining or restoring wild habitats (Navarro and Pereira, 2015; Corlett, 2016), all the while being attentive to WAS and avoiding its increase (Tomasik, 2015). As a consequence, proper metrics of WAS and models supporting decisions to intervene or not are required before any intervention in real ecosystems (Delon and Purves, 2018).

When it comes to decisions related to intervention in ecosystems to reduce WAS, we propose to consider two major positions in this paper. While an Ecological Conservationist (EC) and an Animal Rights Advocate (ARA) both appear to care for animals, they actually pursue objectives that may be at odds with one another, notably because the EC is interested in preserving ecosystems and species, while the ARA is concerned with the suffering of individuals. In 1984, Sagoff showed the fundamental opposition between Peter Singer and Aldo Leopold, a key figure in wilderness conservation (Sagoff, 1984). Peter Singer values equal consideration for the suffering of all individuals while Aldo Leopold values the ecological interactions at the level of ecosystems. Recently, Horta has described some of the disagreements between environmental ethics and concern for wild animal suffering (Horta, 2018). He has outlined that cases for convergence are scarce and uncertain, and suggests that further research may help elicit cases of agreement and disagreement between AR advocates and environmentalists as well as help decide how best to intervene.

In this paper we shall attempt to model the suffering in ecosystems using classical population dynamic models and see how the preferences of an ARA and an EC compare. Roughly, our assumption is that the ARA is interested in minimizing suffering while the EC is interested in maintaining the maximum number of species. In addition to the modelling, we will loosely illustrate the ARA and EC preferences with real cases in different European countries.

## 2 Modeling Wild Animal Suffering : preliminary hypotheses

Let us first describe preliminary hypotheses that are needed in order to justify the use of the population dynamics formalism. Animal sentience can be measured via models or scales, some of which are already used in medicine and behavioral sciences (Paul et al., 2005; Morton et al., 2005; Proctor et al., 2013; Reid et al., 2017; Abboud et al., 2020). However, such scales or models do not exist for wild animals, which makes the modeling of WAS challenging. WAS occurs because wild animals experience various types of harm, such as diseases, injuries, starvation, dehydration, natural disasters, predation, etc. At this stage, modelling the level of suffering associated with all the types of harm is beyond the scope of this article. However, we assume that in most circumstances where such harm occurs, it almost always results in painful deaths. By attributing an equal weight to each harm, we formulate the following hypothesis

(*H*_1_) : *A measure of harms that cause WAS is a measure of deaths occurring among wild animals*.

Of course, *H*_1_ is questionable, because animals living a stressful life do not necessarily die. There are other types of harm that cause suffering without leading to death. Nonetheless, we keep *H*_1_ under the assumption that suffering which does not lead to death is small compared to suffering that does.

Next, we assume that most deaths occur because of harm that has caused suffering. So we formulate the following second hypothesis

(*H*_2_) : *A measure of deaths among wild animals is a measure of WAS*.

Combining *H*_1_ and *H*_2_ enables a convenient metric for measuring WAS: deaths. This idea has already been proposed by Fischer and Lamy in the context of cropping in agriculture (Fischer and Lamey, 2018).

Dynamic population modeling provides a conceptual mathematical framework to describe the dynamics of biological systems that include single species as well as multiple species interactions (Brauer et al., 2012). Since most population variables are time dependent, our metric for suffering is the death rate (mortality rate), symbolized by the variable *m*′ (*t*), where priming refers to differentiation with respect to time. In other words, *m*′ is a measure of the increase in deaths per unit time.

To provide a formal definition on the preferences of the ARA and the conservationist, we consider two hypothetical scenarios *A* and *B*, where the respective total death rates are 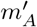 and 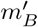, and where the respective total number of species are *N*_*A*_ and *N*_*B*_. On the one hand, since the ARA wants to minimize suffering, according to *H*_1_ and *H*_2_, their preference may be described as follows

**ARA preference** : If 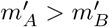and if *N*_*A*_ ≠ 0, then *A* is preferred to *B*. If 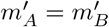, the ARA is indifferent.

We have introduced a condition for non-zero total population because we are assuming the ARA is not an extinctionist, which is a trivial solution for which WAS is rigorously null.

The conservationist wants to maximize the number of species and their preference are described as follows

**Conservationist preference** : if *N*_*A*_ > *N*_*B*_, *A* is preferred to *B*. If *N*_*A*_ = *N*_*B*_, the conservationist is indifferent.

Of course, expressed in this manner, our descriptions may be viewed as exaggerated or incomplete. However, our goal is to reveal compatibilities and oppositions between fundamental preferences. So with the preferences of the EC and the ARA formally stated, we may introduce population dynamic models that enable counting deaths in various situations.

## 3 Population dynamic models

The mathematics of population dynamics is an old field and the following models presented here are very well-known (Brauer et al., 2012). We explore four models and derive the final death rate as a metric of the quantity of suffering for each of them. The details of the models, demonstrations and stability analysis are provided in the supporting information.

### 3.1 Single population competition

Competition models are models that describe individuals that compete for the same resource. The logistic equation models the dynamics of a single population of animals *x*(*t*) that are all in competition for resources in the environment

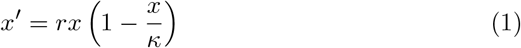

where *r* ∈ ℝ_+_ is the per capita growth rate and *κ* ∈ ℝ_+_ is the carrying capacity. The solution to this equation is analytical

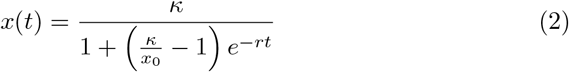

where *x*_0_ = *x*(0) is the initial population. The limit at infinite times, tells us that the size of the population ends up being equal to the carrying capacity at a sufficiently long time. So, aside from total extinction, the equation has a single equilibrium point 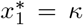, which acts as an attractor (see phase diagram on Fig. 1(a)). If *x*_0_ < *κ* (resp. *x*_0_ > *κ*), the population will grow (resp. decrease) towards *κ* (see Fig. 1(b)).

**Figure 1:**
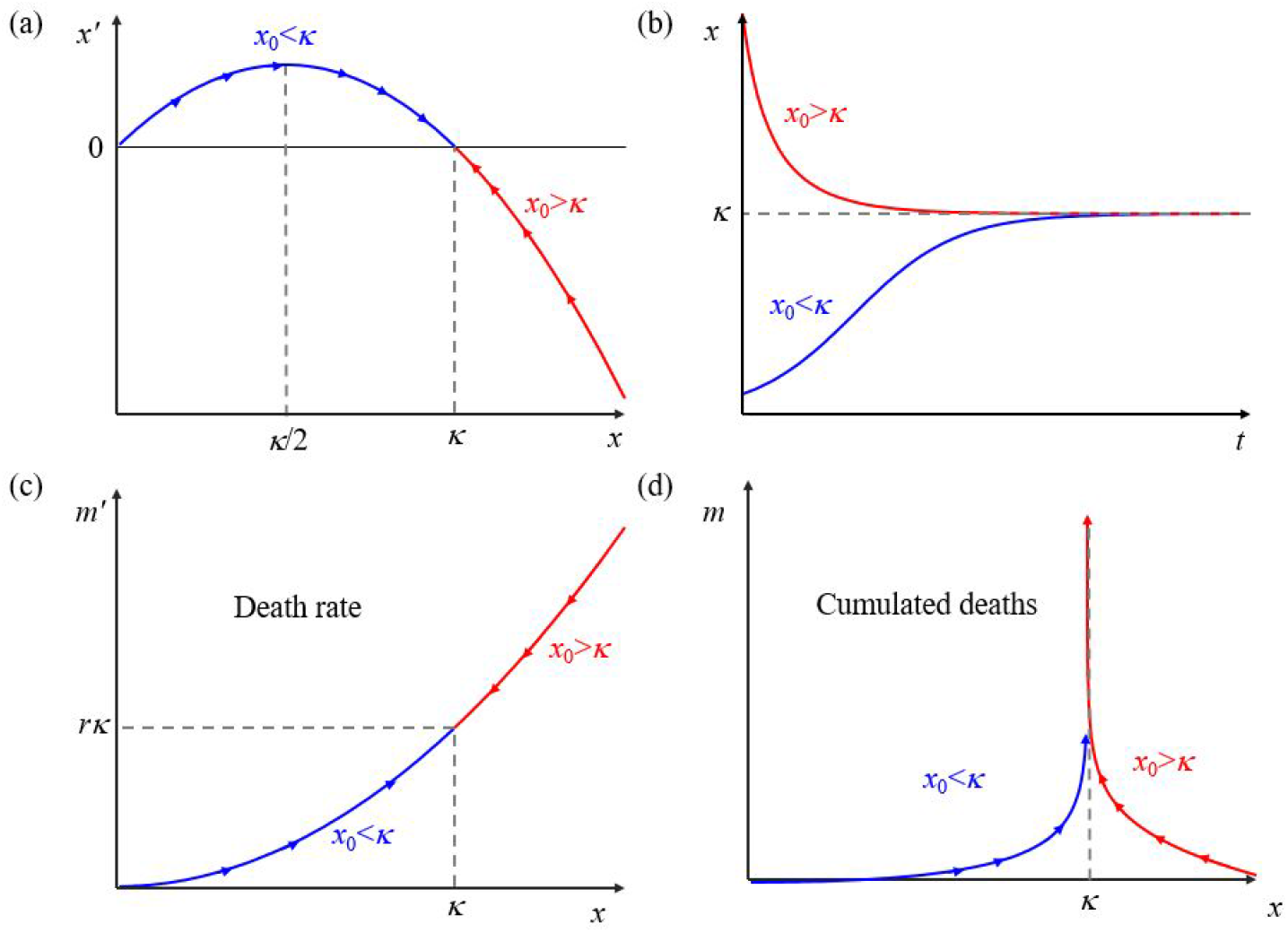
Graphical representations for the single population equations (a) Typical phase diagram of the logistic equation, for *x*_0_ *< κ* (blue stream line) and *x*_0_ *> κ* (red stream line). (b) Two examples of solutions to the logistic equation. The red (blue) curve is obtained for an initial population *x*_0_ that is larger (smaller) than the carrying capacity *κ*. (c) Death rate *m*′ as a function of *x*. (d) Cumulative death rate as a function of *x*.

Figure 1(c) shows the death rate as a function of *x* and is simply

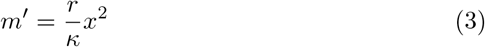

*M*′ cannot be minimized. However we clearly see that as time increases, though the final death rates are identical, the *x*_0_ *< κ* case has a slower increase and as a result minimizes deaths overall (see Fig. 1(d)).

### Two population competition

In a two population competition model, we are concerned with describing the dynamics of two populations over time (*x*(*t*), *y*(*t*)), each of which has its own logistic equation

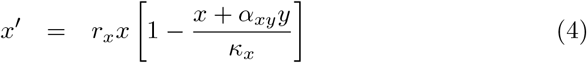

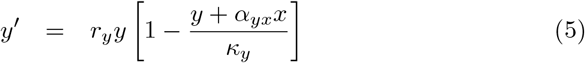

*x* and *y* each have their associated growth rates (*r*_*x*_,*r*_*y*_) and carrying capacities (*κ*_*x*_, *κ*_*y*_). *α*_*xy*_ and *α*_*yx*_ represent the interaction between the two populations. The interaction is competitive (resp. cooperative) if these terms are positive (resp. negative). Here, we only consider strictly positive real values of (*r*_*x*_, *r*_*y*_) and (*α*_*xy*_, *α*_*yx*_). Effectively, this means that the two populations compete and that there are no synergies.

The stability analysis provides four different scenarios that are a function of the values of *a*_*xy*_ = *α*_*xy*_*κ*_*y*_*/κ*_*x*_ and *a*_*yx*_ = *α*_*yx*_*κ*_*x*_*/κ*_*y*_ with respect to 1 (see Fig. 2). The first three cases (see Figs. 2(a-c)) are extinction scenarios, as one population is driven to extinction, and happens when only one coefficient is smaller than 1 or both coefficients are larger than 1. The last is a coexistence scenario (see Fig. 2(d)), which occurs when both coefficients are smaller than 1.

**Figure 2:**
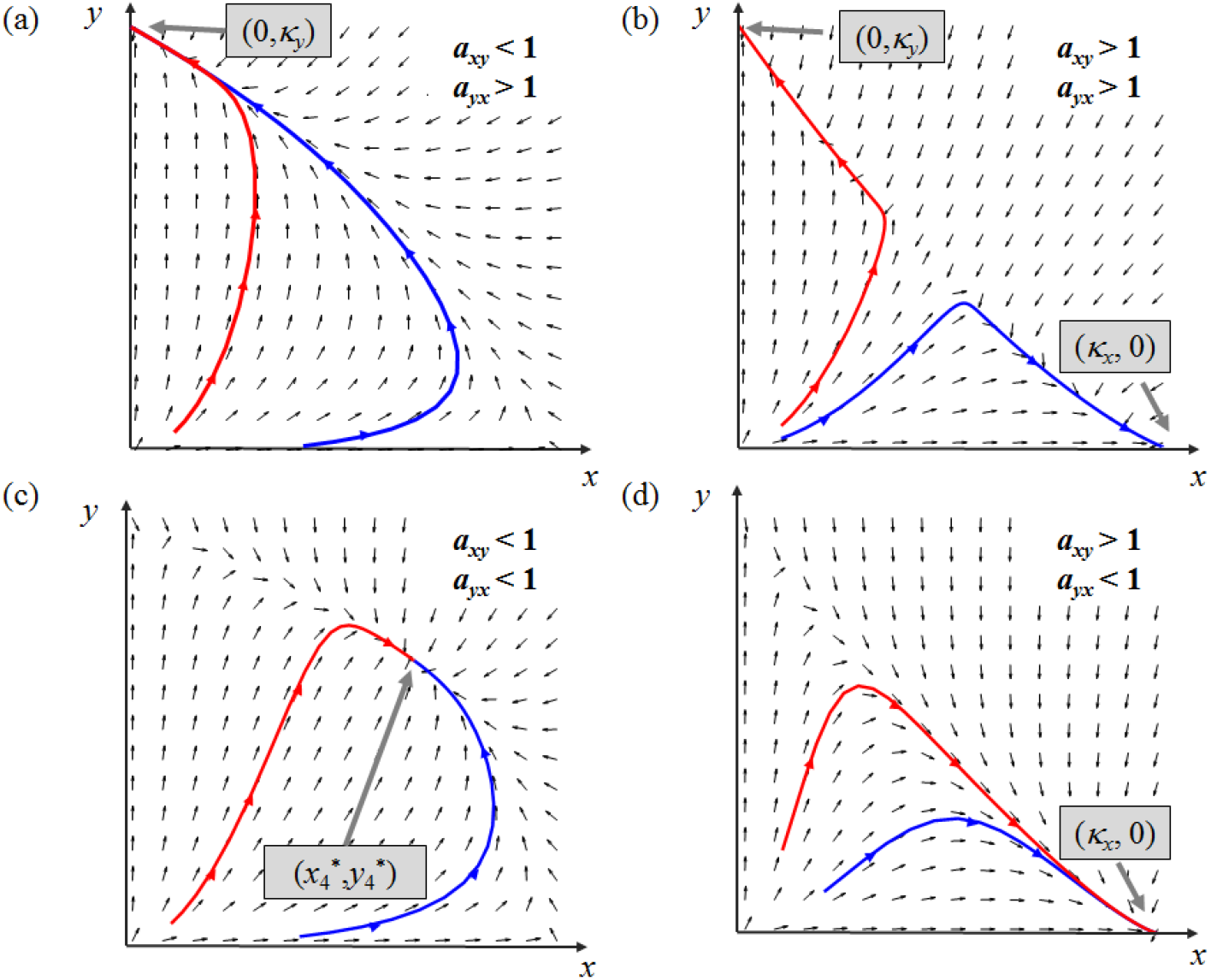
Graphical representations of the two-population solutions to the competitive Lotka-Volterra equations. Each panel is a typical *y* vs *x* phase diagram and the red and blue curves exhibit two possible trajectories that only depend on the initial conditions. The plots are given for the following cases : (a) *a*_*xy*_ < 1 and *a*_*yx*_ > 1, (b) *a*_*xy*_ > 1 and *a*_*yx*_ > 1, (c) *a*_*xy*_ < 1 and *a*_*yx*_ < 1 and (d) *a*_*xy*_ > 1 and *a*_*yx*_ < 1.

The final death rate is equal to 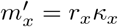 (reps. 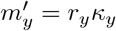), if *y* (resp. *x*) goes extinct in the extinction scenarios. In the coexistence scenario ((*a*_*xy*_, *a*_*xy*_) ∈ [0, 1[^2^), the final death rate can be written in the following compact form

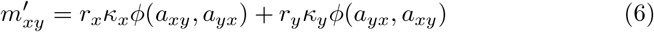

where *ϕ*(*a, b*) = (1 − *a*)*/*(1 − *ab*).

### The classical Predator-Prey model

In the predator-prey model, also known as the Lotka-Volterra equations, there is a prey population *x*, which is assumed to exhibit exponential growth in the absence of predators and a predator population *y*, assumed to exhibit exponential decay in the absence of prey

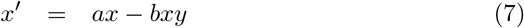

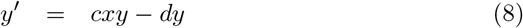

where *a* is the prey growth rate, *b* is the death rate per unit predator. *c* is the growth rate per unit prey and *d* is the predator death rate. In this model, prey only die because of predation and predators can only grow by predation. If all coefficients (*a,b,c,d*)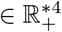 and excluding the trivial total extinction point, the only stable equilibrium is 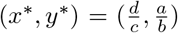, where solutions form closed orbits around this equilibrium point. The amplitude of the orbit depends on the initial conditions. A stable orbit necessarily exists as long as the initial conditions *x*_0_ and *y*_0_ are simultaneously non-zero. Figure 3 shows two typical trajectories in the *y* vs *x* phase diagram that form orbits around 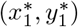. The death rate is expressed as

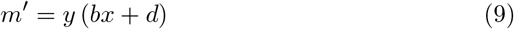

The death rate can be minimized if the predator population is zero (*y* = 0), but at the expense of unlimited exponential growth of the prey population, which is unrealistic, or may be used only in situations where the prey population is very far from the environmental carrying capacity.

**Figure 3:**
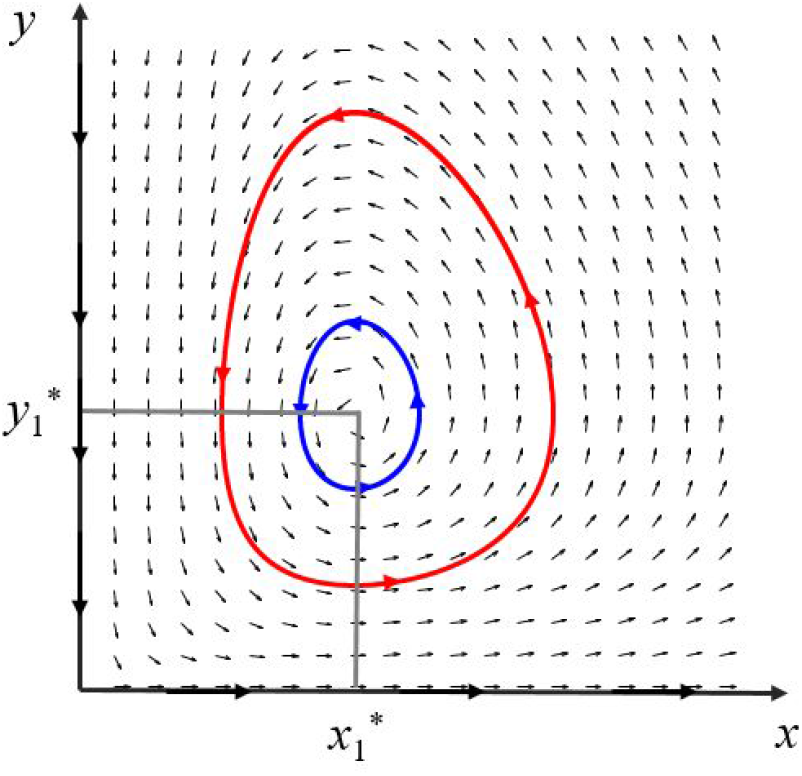
Phase diagram (*y* vs *x*) of the predator prey Lotka-Volterra model. Two typical orbits are represented. 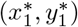 is the equilibrium point around which stable orbits are centered.

### The Predator-Prey model with a carrying capacity for prey

To circumvent limitless exponential growth of the prey population in the absence of a predator, a carrying capacity term *κ* for the prey can be introduced.

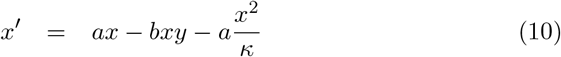

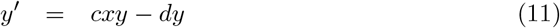

There are two equilibrium points aside from total extinction. The first is 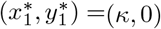, which happens if 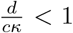. It is a saddle reachable from the x-axis when *y* = 0. If 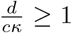, the point is a stable sink. Predators go extinct and the prey reach their carrying capacity. In this case, similarly to the single population competitive case, the final death rate is 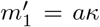. The second equilibrium is 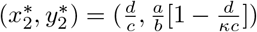, which can only exist under the condition that 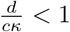. The point is a sink and corresponds to a stable coexistence scenario. In this case, the final death rate is

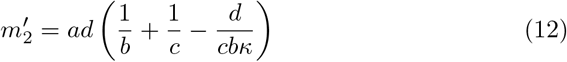

## 4 Animal right’s advocacy vs. conservationism

In this section we evaluate the different models and possible scenarios based on the positions of an ARA and an EC. As a reminder, the ARA is not an extinctionist, meaning that his aim is not to have all species go extinct to minimize suffering. As a result, they might agree to let one species go extinct in a two-species situation, but they will never agree to having both species go extinct. This means that we should specify what the decision variables are. The decision variables are those for which the decision-makers (here the ARA and EC) have a certain degree of control.

Figure 4 below summarizes the different scenarios, different examples from practice, relevant decision variables, final death rates and preferences of the ARA and EC. The following paragraphs detail, for each type of model, the agreement or disagreement between each position depending on ecosystems parameters.

**Figure 4:**
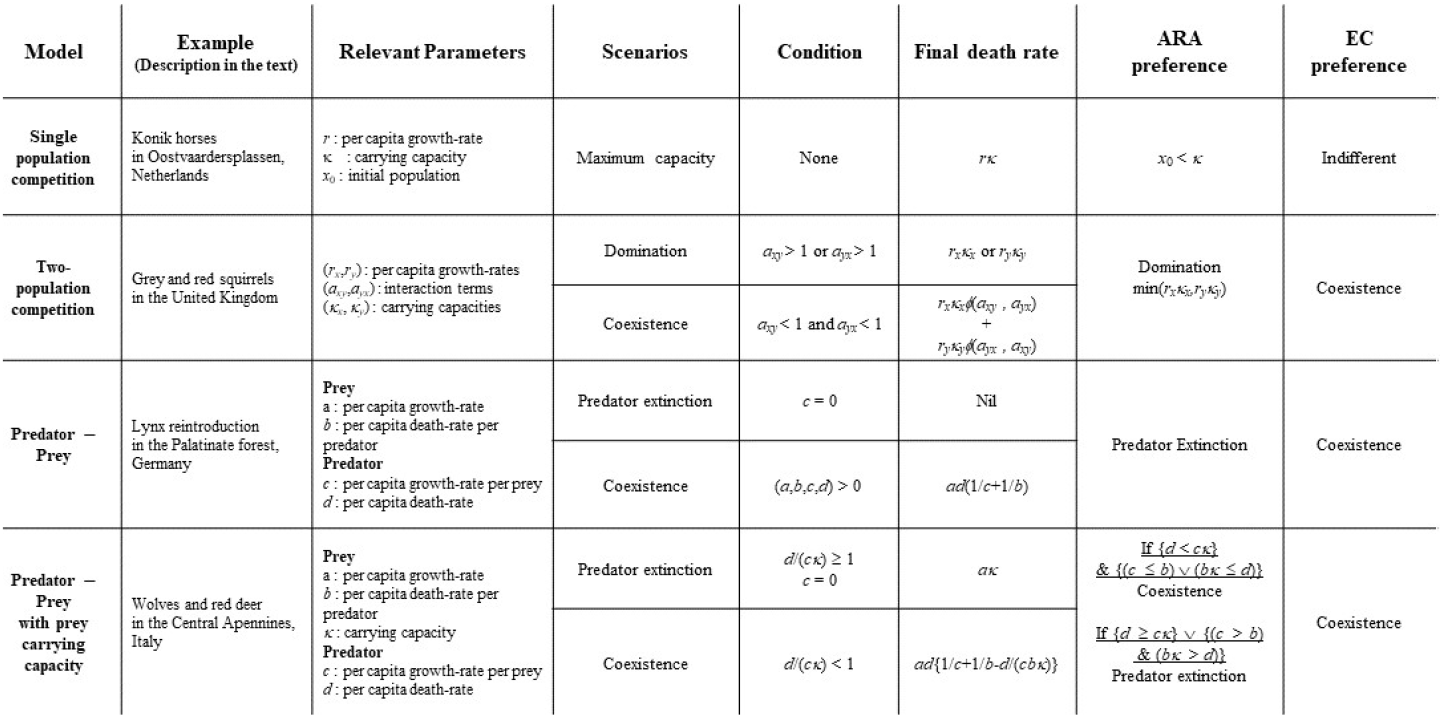
Table summarizing for each model and example from practice, the different equilibrium scenarios, relevant decision parameters, conditions to reach that scenario, final death and preferences of the animal rights advocate (ARA) and ecological conservationist (EC).

Overall, we consider the decision variables to be the parameters that control growth-rates, death-rates, interaction terms and carrying capacities. For instance, vaccinating a given species is likely to increase its growth-rate or interaction term. Aside from this example, we shall not describe how concrete measures may influence each variable.

### 4.1 Single population competition

In the single population competition model, the final death rate is *m*^′^ = *rκ*. An EC would most probably be indifferent to the model decision variables since the equilibrium population corresponding to the carrying capacity is always reached. An ARA on the other hand wants to minimize WAS, which in the context of this model means minimizing the overall deaths. This can be done by approaching *κ* from lower values, which means making sure *κ > x*_0_. Interestingly, we see that the growth-rate is an irrelevant decision variable. As a result, the EC and ARA can be brought into agreement for a single population competition model. For this, consider the “wildlife experiment” (Lorimer and Driessen, 2014) in the Oostvaarderplassen nature reserve in the Netherlands. In the 1980s, large herbivores such as cattle, horses, and red deer corresponding to a Pleistocene grazing guild were introduced to increase vegetation heterogeneity. Heck cattle and Konik horses replace the extinct Aurox and Tarpan (wild forest horse). Management within the 60 km^2^ fenced area is limited to compliance with basic animal welfare standards. Populations are regulated only density-dependent or by interspecific competition, i.e., many animals may starve to death during harsher winters (Vera, 2009). Considering only intraspecific competition, ARAs would be against the approach if *x*_0_ *> κ*, e.g. if herbivores appear malnourished or even die, while ECs would accept these conditions.

### 4.2 Two populations competition

In the two-population-competition model, the EC would favor coexistence since they are interested in maximizing the number of species preserved. To minimize WAS, the ARA would only accept this choice if the final death rate in the coexistence scenario were smaller than any of the two death rates involved in the domination scenarios. This can be summarized by the following inequality

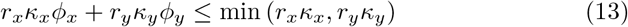

where *ϕ*_*x*_ = *ϕ*(*a*_*xy*_, *a*_*yx*_) and *ϕ*_*y*_ = *ϕ*(*a*_*yx*_, *a*_*xy*_). However, this condition can never be reached for (*a*_*xy*_, *a*_*xy*_) ∈]0, 1[^2^, because *ϕ*_*x*_ + *ϕ*_*y*_ is always greater than 1. As a consequence, the ARA shall choose to keep the species that has the minimal death rate and let the other be driven to extinction. In the two-population competition model, the conservationist and ARA have incompatible preferences. As a consequence, the ARA shall choose to keep the species that has the minimal death rate and let the other be driven to extinction. So it seems that for the two-population competition model, the EC and ARA have incompatible preferences. Here we may look again at the case of the Oostvaarderplaassen reserve and now consider the interspecific competition between the different herbivore species. ARAs would aim for managing the guild by only keeping the species with the lowest death rate, whereas ECs would let all species coexist. Another example is the competition of native red squirrels with gray squirrels, introduced from the America to the UK in the late 19th century (Okubo et al., 1989) and now spreading across Europe (Bertolino et al., 2008). Competition over food resources and a virus transmitted by gray squirrels as vector species is causing red squirrel populations to decline in many European areas (Strauss et al., 2012). According to an ARA, the appearing dominance of gray squirrels is justified due to their lower mortality rate compared to red squirrels. In contrast, an EC would favor coexistence of the two species, implying that action would be needed to reduce gray squirrel populations and ensure the survival of red squirrels. It should be noted here, that alternate conservationist approaches may even advocate for the removal of the grey squirrel population in order to revert to the past situation prior to their introduction through human intervention. For the sake of simplicity, we have not considered such approaches.

### 4.3 Predator-prey

In the classical predator-prey model, the EC and ARA would disagree on the strategy as well. While the EC will be interested in (at least) reaching a stable orbit around any equilibrium point, they require simultaneous non-zero populations for both the prey and predator. The ARA would want to drive the predators to extinction, so that the death rate can be zero. As mentioned previously, extinction is not a very plausible scenario, because of limitless exponential growth of the prey population. To this end, we can take a closer look at a lynx reintroduction project in the German-French UNESCO Biosphere Reserve Palatinate forest/Northern Vosges. Between 2016 and 2020, a total of 20 lynx from the Slovakian Carpathians and Switzerland were reintroduced to establish a subpopulation and restore predation in the ecosystem (SNU). Their main prey, roe deer, finds a well-suited habitat in the large forest ecosystem with year-round mild weather conditions. We may therefore assume a high carrying capacity of the roe deer population and neglect this variable in the present case. Here, ARAs would argue against the introduction of a predator into the ecosystem, while ECs would favor coexistence of lynx and deer.

### 4.4 Predator-prey with carrying capacity

Finally, for the predator-prey model with a carrying capacity, comparing the death rates in the extinction of predators or predator-prey coexistence scenarios, it is useful to consider the following polynomial function constructed with the ratio of 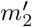 to 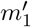

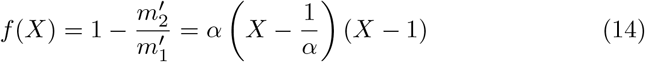

where *α* = *c/b* and *X* = *d/*(*cκ*). For the ARA to accept the coexistence scenario, they would require *f* (*X*) ≥ 0. *f* has two roots : *X*_1_ = 1*/α* and *X*_2_ = 1. The coexistence scenario only occurs for *X* ∈] 0, 1[, so there are two possible situations depending on the value of *α*. If *α* ≤ 1, then both roots are greater or equal to 1, and since *f* is convex it is positive for all *x <* 1 so the ARA prefers coexistence. If *α* ≥ 1, then *X*_1_] 0, 1 [and since *f* is convex, it is positive for all *X* ≤ 1*/α*. In summary, if a coexistence scenario is possible, that is *X <* 1, then the ARA shall agree to coexistence if either of these two conditions are met

1. *bκ* ≤ *d*
2. *c* ≤ *b*

We see that in these two situations the ARA and EC can agree to the coexistence scenario. *bκ/d* ≤ 1 may be viewed as a sufficient condition for the ARA and the EC to agree to coexistence, but not a necessary one. In plain words, for the ARA and the EC to agree, we need the following conditions to be met:

i. **Coexistence condition** : The predator per capita death-rate should be smaller than the predator per capita growth-rate had the prey population reached its carrying capacity (*d < cκ*).
ii. **Agreement condition** : The predator growth-rate per unit prey should be less or equal to the prey death-rate per unit predator (*c* ≤ *b*) or the prey death-rate per unit predator, had the prey population reached its carrying capacity, should be less or equal to the predator death-rate (*bκ* ≤ *d*).

As an example, we want to take a closer look at wolf-red deer dynamics in the Central Apennines, a mountain range in the Italian peninsula. Both Apennine wolf and red deer populations live in high densities in this region (Rewilding Apennines, 2021). During snowy and cold winters, red deer compete for scarce food resources, so we can assume that they approach their carrying capacity under these conditions. Assuming very harsh winters, ARAs and ECs would agree that these species coexist as long as over the course of the year, the per capita growth rate of wolves per unit red deer is less than the per capita death rate of red deer per unit wolf. Indeed, we can observe much higher numbers of red deer than of wolves in the Central Apennines, which are probably following the typical predator-prey power laws of terrestrial trophic cascades (Hatton et al., 2015). Thus, in this case, we would find an agreement between ARAs and ECs in terms of arguing for wolf and red deer coexistence.

## 5 Perspectives to improve the modeling of WAS

### 5.1 A wider range of human perspectives

Our results show that in some situations with regards to the type of ecosystem, humans interested in reducing WAS and those interested in species conservation can have compatible preferences and intervene in favor of WAS reduction while promoting biodiversity. These results comfort previous views that conservation and consideration for animal welfare could cooperate (Fraser, 2010). This is far from obvious since cases of conflicts have been documented between conservation objectives and animal ethics (Jones et al., 2012). As a matter of fact, the traditional view is that animal ethics and conservation are incompatible (Hutchins and Wemmer, 1987). A central issue is that animal ethics focuses on caring for individuals which is at odds with the conservationist position of caring for species. Some authors promote “compassionate conservation” as an attempt of getting the care for individuals to comply with the care for ensembles of individuals (species, populations, etc.), resulting in an attempt to limit harm in conservation (Bekoff, 2013; Wallach et al., 2018). However this proposal is challenged by some authors who consider that empathy spoils rational utilitarian decision-making, which should ultimately drive conservation for the long-term survival of our planet’s ecosystems (Griffin et al., 2020). Other authors argue that, while compassion is a laudable objective, it should not be at the expense of the survival of species or their habitat or even of human life safeguarding (Oommen et al., 2019).

Additionally, our model merely approaches WAS from a utilitarian perspective and does not adopt other philosophical perspectives on the topic. For instance, both the ARA and conservationist can also root their action on deontological principles (Macdonald et al., 2016; Regan, 2004). This means that even if utilitarian ARAs and conservationists were to agree and cooperate, this cooperation could be challenged by other non-consequential views on the matter. Furthermore, in reality ARA and EC preferences are probably not only determined by mortality and number of wild animal populations. For example, ARAs might additionally account for the suffering of livestock from attacks by wild predators when considering predator-wild prey relationships. ECs might value the suffering of an endangered species to be more severe than that of a non-endangered species. Finally, our definition of an ecological conservationist as being a person interested in maximizing the number of species may seem a little too restrictive and falls under an ecocentric form of environmentalism, meaning maximizing interactions in ecosystems. Indeed, there exists other forms of environmentalism that may have alternative preferences such as naturocentric and biocentric (Horta, 2018). A naturocentric view prefers ecosystems without human interference. A biocentric view gives priority to the value of individual life forms. While biocentric environmentalism, close to compassionate conservation, may very well have preferences that are more compatible with the ARA preferences, it would be less likely for naturocentric conservationists. Overall, our simple approach is a first step and certainly needs some refinement to account for all the various forms of environmentalism and animal ethics.

### 5.2 Improving WAS modeling

The first element that could be improved further is the modeling of the suffering. Suffering is a multidimensional problem that can integrate many factors such as hunger, stresses caused by predation, disease, wounds, cold, heat, effects of natural hazards and the impossibility for a wild animal of expressing normal behavior. The comparison of wild animal behavior with the one living in captivity can give cues on the latter but can not be relied upon as an objective measure (Veasey et al., 1996). Integrating these complex factors will certainly enhance the WAS picture but research is still lacking to give a precise method to asses WAS. Due to this limitation and in spite of large amounts of suffering in the wild, our results suggest that we are still very far from having proper models to legitimize any intervention. There is a significant need for further empirical studies to find an objective assessment of WAS which would improve its modeling and evaluate the potential of intervention. In that sense, national parks and pro-wilderness programs have a strong role to play in increasing our knowledge of WAS and the consequences of intervention in attempting to reduce it.

A second element that could improve the modeling of WAS is a recent approach in ecological modeling based on the individual rather than the group (Stillman et al., 2015). Because suffering is an experience felt at the individual level, this agent-centered approach could focus on the set of experiences and animal behavior that can lead an individual to suffer. This approach would require the research and elaboration of detailed models of behavior for each type of animal in a studied ecosystem (see for example (Duriez et al., 2009; Carter et al., 2015)). Such individual models could then be integrated into an agent-based model that can simulate interactions between agents (like predation for example) (Railsback and Grimm, 2019). Additionally, these models can represent socio-ecological systems and thus have human agents interacting with wild species (Le Page et al., 2015). Due to the possibility of having different types of human agents, such models could include diverse views among ARAs and conservationists.

## 6 Conclusion

Many scholars agree that intervening in favor of reducing WAS requires a proper understanding of the consequences. This calls for models capable of informing such decisions. We cast a first stepping stone in that direction by using death rates as a metric in the frame of classical population dynamics models. We show that in some situations, with regards to the type of ecosystem, humans interested in reducing WAS and those interested in species conservation can have compatible preferences and intervene in favor of WAS reduction while promoting biodiversity. In other situations, they will disagree because WAS reduction entails driving a species to extinction, which is incompatible with conservationism. This last point shows that as concern for animal welfare grows, the disagreement between animal rights advocates and conservationists may grow as well. Our results invite both actors to first focus on ecosystems where both worldviews can find common ground to reduce WAS while achieving conservation objectives, consistently with Horta’s prescription (Horta, 2018).

## Supporting information

Supplemental information for main manuscript

## Acknowledgments

We would like to warmly thank Dr. Ana Stritih and Dr. Olivier Cailloux for their precious support, feedbacks and advice to improve this article.

## Statements and Declarations

All authors declare that they have no conflicts of interest nor do they have competing interests.

