## Supplemental information for main manuscript for "Conservation vs. Wild-Animal Suffering : how can population dynamics help?"

This document serves as supporting information for our main manuscript and provides additional mathematical detail to the claims made. Section 1 describes the population dynamic models in more depth. Section justifies the preference incompatibility of the ARA and the EC in the two-population competition model. Section 3 justifies the cases for agreement between the ARA and EC in the predator-prey model with a carrying capacity for prey. Finally, section 4 provides details on the method used to analyse the stability of equilibrium points and applies it to each model considered in the main manuscript.

#### 1 Population dynamic models

The mathematics of population dynamics is an old field and the following models presented here are very well-known. We present them and try to be as pedagogical as possible, but we do not claim to bring any mathematical novelty into these models, nor do we claim to be comprehensive. We direct the interested reader towards Brauer and Castillo-Chavez' monograph for further information on these models [1]. We are merely interested in deriving the final death rate for each model considered and establish it as a metric of the quantity of suffering. This will then serve to determine the preferences of an ARA interested in minimizing WAS. In the last section we discuss the preferences of the ARA and compare it to an EC's assumed preferences. The stability analyses for each model is given in section 4.

##### Single population competition

Competition models are models that describe individuals that compete for the same resource. We will start with the logistic equation, which is perhaps one of the simplest and most famous models.

*
†

The logistic equation models the dynamics of a single population of animals $x(t)$ that are all in competition for resources in the environment

$$x' = rx\left(1 - \frac{x}{\kappa}\right) \tag{1}$$

where $r \in \mathbb{R}_+$ is the per capita growth rate and $\kappa \in \mathbb{R}_+$ is the carrying capacity. In this differential equation, priming denotes the mathematical operation of deriving with respect to time. The solution to this equation is analytical

$$x(t) = \frac{\kappa}{1 + \left(\frac{\kappa}{x_0} - 1\right) e^{-rt}} \tag{2}$$

where $x_0 = x(0)$ is the initial population. We see that

$$\lim_{t \to +\infty} x(t) = \kappa \tag{3}$$

and from this, it is obvious that the carrying capacity $\kappa$ represents the maximum population size that is sustainable by the environment. Finding the limit at infinite times, tells us what the population ends up being equal to after a sufficiently long time. This model is simple enough that an analytic solution can be found. For more complicated models, this is not always possible, but in most cases we can still provide strong and true claims about the model even without knowing the solution. The standard procedure consists in finding the equilibrium points, that is, the values of $x$ that cancel $x'$. We call these *equilibrium points*, because when $x' = 0$, $x$ is a constant solution. Equilibrium points are not necessarily stable. Stable points act as attractors, which means that the population will eventually fall towards that point, while unstable points on the contrary act as sources, which means that the population has a tendency to move away from that point. Here, there are two equilibrium points: $x_1^* = 0$ and $x_2^* = \kappa$. $x_1^*$ is unstable, unless the initial population is rigorously null ($x_0 = 0$), while $x_2^* = \kappa$ is an attractor. There are therefore two possible initial states

1. $\underline{x_0 < \kappa}$ : when the initial populations is smaller than the carrying capacity and as a result, $x' > 0$ and $x(t)$ will grow to $\kappa$.

2. $\underline{x_0 > \kappa}$ : when the initial population is larger than the carrying capacity and as result, $x' < 0$ and $x(t)$ will decrease to $\kappa$.

$x_2^* = \kappa$ *attracts* the population towards it. The death rate is the number of deaths per unit time and is simply

$$m' = \frac{r}{\kappa} x^2 \tag{4}$$

It corresponds to the term that is substracted in eq. (3). It is proportional to the square of the population at any time. As a result, there is no way of minimizing it (except for the trivial solution $x = 0$). When the equilibrium is reached, $m' = r\kappa$ and deaths just keep increasing linearly with time : $m(t) = r\kappa t$. As times increases, though the final death rates are identical reagardless of whether $x_0 < \kappa$ or $x_0 > \kappa$, the $x_0 < \kappa$ case has a slower increase and as a result minimizes deaths overall.

#### Two population competition

In a two population competition model, we are concerned with describing the dynamics over time of two populations ($x(t)$ and $y(t)$), each of which has its own logistic equation

$$x' = r_x x \left[1 - \frac{x + \alpha_{xy} y}{\kappa_x}\right] \quad (5)$$

$$y' = r_y y \left[1 - \frac{y + \alpha_{yx} x}{\kappa_y}\right] \quad (6)$$

$x$ and $y$ each have their associated growth rates ($r_x$,$r_y$) and carrying capacities ($\kappa_x$, $\kappa_y$). We have introduced two terms, $\alpha_{xy}$ and $\alpha_{yx}$, which represent the interaction between the two populations. If $\alpha_{xy} > 0$, then the larger $y$, the more this term will have a tendency to decrease the population $x$. This would represent competition. On the contrary if $\alpha_{xy} < 0$, the interaction will have a a tendency to increase the population $x$. This would be cooperation. Applying the change of variables $X = x/\kappa_x$ and $Y = y/\kappa_y$, as well as setting $a_{xy} = \alpha_{xy}\kappa_y/\kappa_x$ and $a_{yx} = \alpha_{yx}\kappa_x/\kappa_y$, the system of differential equations takes the following simpler form

$$X' = r_x X (1 - X - a_{xy} Y) \quad (7)$$

$$Y' = r_y Y (1 - Y - a_{yx} X) \quad (8)$$

Instead of considering the total populations $x$ and $y$, in this system, we are considering the respective fractions of carrying capacity[1]. It this section, we will only be concerned with strictly positive real values of $(r_x, r_y)$ and $(a_{xy}, a_{yx})$. Effectively, this means that the two populations compete and that there are no synergies.

In summary, the stability analyses can be sorted into the following cases that are a function of the values of $a_{xy}$ and $a_{yx}$ with respect to 1. The first two cases are domination scenarios, while the last is a coexistence scenario

- $a_{xy} > 1$ and $a_{yx} < 1$, then $y$ is driven to extinction, while $x$ thrives and reaches its carrying capacity $\kappa_x$. Here the final death rate is : $m'_x = r_x\kappa_x$.
- $a_{xy} < 1$ and $a_{yx} > 1$, then $x$ is driven to extinction, while $y$ thrives and reaches its carrying capacity $\kappa_y$. Here the final death rate is : $m'_y = r_y\kappa_y$
- $a_{xy} > 1$ and $a_{yx} > 1$, then both populations compete and one wins, while the other is driven to extinction. Either population can win depending on the initial conditions. Here the final death rate is either $m'_x$ or $m'_y$ depending on which population thrives.
- $a_{xy} < 1$ and $a_{yx} < 1$, then a stable equilibrium is reached where both populations coexist

$$x_4^* = \kappa_x \left(\frac{1 - a_{xy}}{1 - a_{xy}a_{yx}}\right) \quad (9)$$

$$y_4^* = \kappa_y \left(\frac{1 - a_{yx}}{1 - a_{xy}a_{yx}}\right) \quad (10)$$

[1]For instance, considering $X = 1$ means that the population is at its carrying capacity. If $X = 2$, then the population is at twice its carrying capacity. This should be taken into account if we are to consider numbers of individuals rather than fractions of carrying capacity.

Here the final death rate is

$$m'_{xy} = m'_x \frac{1 - a_{xy}}{(1 - a_{xy}a_{yx})} + m'_y \frac{1 - a_{yx}}{(1 - a_{xy}a_{yx})} \tag{11}$$

which can be rewritten in the following compact form

$$m'_{xy} = r_x \kappa_x \phi(a_{xy}, a_{yx}) + r_y \kappa_y \phi(a_{yx}, a_{xy}) \tag{12}$$

#### The classical Predator-Prey model

In the predator-prey model, also known as the Lotka-Volterra equations, there is a prey population $x$, which is assumed to exhibit exponential growth in the absence of predators and a predator population $y$, assumed to exhibit exponential decay in the absence of prey

$$x' = ax - bxy \tag{13}$$
$$y' = cxy - dy \tag{14}$$

There is an interaction term ($bxy$), which acts a sink for preys and another ($cxy$), which acts as a source for predators. $a$ is the prey growth rate, while $b$ is the death rate per unit predator. $c$ is the growth rate per unit prey and $d$ is the predator death rate. In this model, prey only die from being *eaten* by predators and predators can only grow by *eating* prey. If all coefficients $(a, b, c, d) \in \mathbb{R}_+^{*4}$, there are three possible equilibrium points

1. $\underline{(x_0^*, y_0^*) = (0, 0)}$ : This point is a saddle, that can only be reached if $x = 0$ and predators wil gradually go extinct from starvation.

2. $\underline{(x_1^*, y_1^*) = (\frac{d}{c}, \frac{a}{b})}$ : solutions form closed orbits around this equilibrium point. The amplitude of the orbit depends on the initial conditions. A stable orbit necessarily exists as long as the initial conditions $x_0$ and $y_0$ are simultaneously non-zero.

The death rate is expressed as

$$m' = y\,(bx + d) \tag{15}$$

This is an actual situation where the death rate can be canceled without driving both populations to extinction. Indeed, this happens, when the predator population is null. Then the prey population exponentially grows. As a result, under the initial condition $x_0 \neq 0$ and $y_0 = 0$, the prey dynamics have the following solution

$$x(t) = x_0 e^{at} \tag{16}$$

Of course, this population growth is explosive and unrealistic since there is always a carrying capacity to the environment.

#### The Predator-Prey model with a carrying capacity for prey

The classical predator-prey model is not very realistic because it permits limitless exponential growth of the prey population in the absence of a predator. Of course, it is implausible that any environment can withstand an infinite population. To circumvent this situation, we can introduce a carrying capacity term $\kappa$ for the prey, as in the single population model

$$x' = ax - bxy - a\frac{x^2}{\kappa} \quad (17)$$

$$y' = cxy - dy \quad (18)$$

There are three equilibrium points

1. $\underline{(x_0^*, y_0^*) = (0,0)}$ : this point is a saddle that can only be approached from the y-axis if $x = 0$.

2. $\underline{(x_1^*, y_1^*) = (\kappa, 0)}$ : If $\frac{d}{c\kappa} < 1$, then it is a saddle point that can only be reached from the x-axis when $y = 0$. If $\frac{d}{c\kappa} \geq 1$,then this point is a stable sink. This is a case of course, where the predators would go extinct and the prey reach their carrying capacity. In this case, similarly to the single population competitive case, the final death rate is $m_1' = a\kappa$

3. $\underline{(x_2^*, y_2^*) = (\frac{d}{c}, \frac{a}{b}[1 - \frac{d}{\kappa c}])}$: of course, since there are no negative populations, this point can only exist under the condition that $\frac{d}{c\kappa} < 1$, which is the saddle point condition of $(x_1^*, y_1^*)$. This point is a sink and corresponds to a stable coexistence scenario. Here the final death rate is

$$m_2' = ad\left(\frac{1}{b} + \frac{1}{c} - \frac{d}{cb\kappa}\right) \quad (19)$$

### 2 Justification for preference incompatibility in the two-population competition model

As stated in the main manuscript, in the two-population-competition model, the EC and ARA have incompatible preferences. Here is a detailed justification of why this is true. To minimize WAS, the ARA would only accept keep if the final death rate in the coexistence scenario were smaller than any of the two death rates involved in the domination scenarios. This can be summarized by the following inequality

$$r_x\kappa_x\phi_x + r_y\kappa_y\phi_y \leq \min\left(r_x\kappa_x, r_y\kappa_y\right) \quad (20)$$

where $\phi_x = \phi(a_{xy}, a_{yx})$ and $\phi_y = \phi(a_{yx}, a_{xy})$. If we choose to label $x$ the population with the lowest final death rate, then this last equation can be divided by $m_x' = r_x\kappa_x$ which yields

$$\phi_x + \rho\phi_y \leq 1 \quad (21)$$

where $\rho = r_x\kappa_x/(r_y\kappa_y) \geq 1$.

Since 1 is an upper bound for $\rho$, a sufficient condition for the previous inequality to be violated is that $\phi_x + \phi_y > 1$. Let us rewrite $f(a_{xy}, a_{yx}) = \phi_x + \phi_y$, by applying the change of variable $(a_{xy}, a_{yx}) = (1 - \varepsilon_x, 1 - \varepsilon_y)$, where $(\varepsilon_x, \varepsilon_y) \in ]0,1[^2$. We then see that

$$f(1 - \varepsilon_x, 1 - \varepsilon_y) = \frac{\varepsilon_x + \varepsilon_y}{\varepsilon_x + \varepsilon_y - \varepsilon_x\varepsilon_y} > 1 \quad (22)$$

which violates eq.(21). As a result, The ARA and EC have incompatible preferences.

### 3 Justification of cases for the predator-prey model with carrying capacity for prey

In the main manuscript, we detail the conditions for agreement between the ARA and EC in the predator-prey model with a carrying capacity for prey. A justification of these conditions is given here.

For the predator-prey model with a carrying capacity, comparing the death rates in the extinction of predators or predator-prey coexistence scenarios, it is useful to consider the following polynomial function constructed witht the ratio of $m'_2$ to $m'_1$

$$f(X) = 1 - \frac{m'_2}{m'_1} = \alpha \left( X - \frac{1}{\alpha} \right) (X - 1) \tag{23}$$

where $\alpha = c/b$ and $X = d/(c\kappa)$. For the ARA to accept the coexistence scenario, they would require $f(X) \geq 0$. $f$ has two roots : $X_1 = 1/\alpha$ and $X_2 = 1$. The coexistence scenario only occurs for $X \in ]0,1[$, so there are two possible situations depending on the value of $\alpha$. If $\alpha \leq 1$, then both roots are greater or equal to 1, and since $f$ is convex it is positive for all $x < 1$ so the ARA prefers coexistence. If $\alpha \geq 1$, then $X_1 \in ]0,1[$ and since $f$ is convex, it is positive for all $X \leq 1/\alpha$. In summary, if a coexistence scenario is possible, that is $X < 1$, then the ARA shall agree to coexistence if either of these two conditions are met

1. $b\kappa \leq d$

2. $c \leq b$

We see that in these two situations the ARA and EC can agree to the coexistence scenario. $b\kappa/d \leq 1$ may be viewed as a sufficient condition for the ARA and the EC to agree to coexistence, but not a necessary one. In plain words, for the ARA and the EC to agree, we need the following conditions to be met:

i/ **Coexistence condition** : The predator per capita death-rate should be smaller than the predator per capita growth-rate had the prey population reached its carrying capacity ($d < c\kappa$).

ii/ **Agreement condition** : The predator growth-rate per unit prey should be less or equal to the prey death-rate per unit predator ($c \leq b$) or the prey death-rate per unit predator, had the prey population reached its carrying capacity, should be less or equal to the predator death-rate ($b\kappa \leq d$).

### 4 Detailed stability analysis of equilibrium points for each model

This section is devoted to presenting the method used to analyse the stability of equilibrium points and detail this analysis for every model considered.

#### 4.1 Stability analysis of equilibrium points

Figure S1 is a Poincaré diagram of the equilibrium points for a linear differential system $x' = Ax$. In our study, we linearize the nonlinear differential equations close to the equilibrium points and study the Jacobien $J$. This can be done because making the change of variables $\hat{X} = (X - X^*)$ and $\hat{Y} = (Y - Y^*)$, the following differential system can be produced that studies how small deviations behave near the equilibrium point $(X^*, Y*)$

$$\begin{bmatrix} \hat{X}' \\ \hat{Y}' \end{bmatrix} = J \begin{bmatrix} \hat{X} \\ \hat{Y} \end{bmatrix} \tag{24}$$

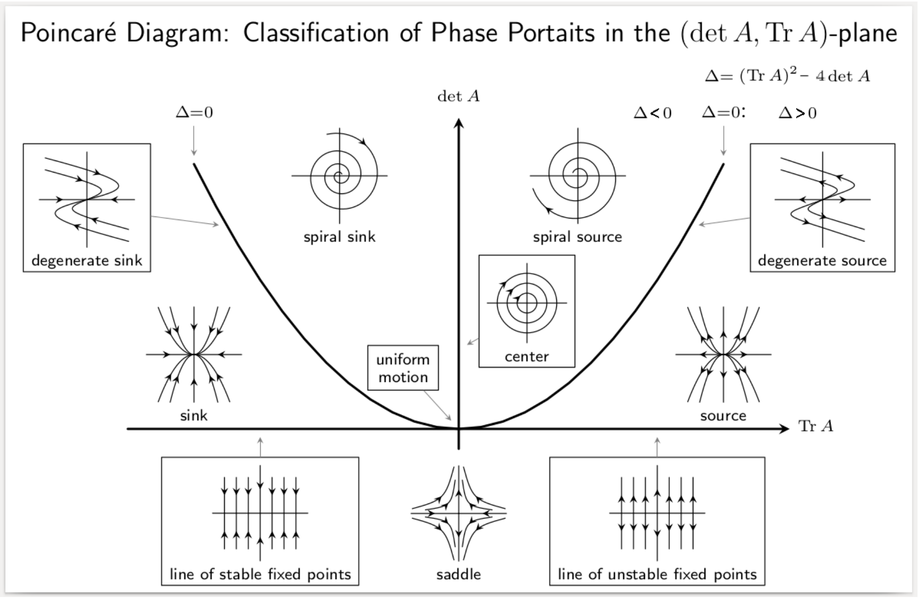


Figure S1: Equilibrium point stability diagram for a linear autonomous system $x' = Ax$. This diagram may be used to classify equilibriums according to their features (adapted from wikipedia).

#### 4.2 Two-population competition

In what follows, we shall determine the equilibrium points $(X^*, Y^*)$ in the frame of the two-population competition model described in the main text and then analyze their stability by linearizing the differential system near these points by using the Jacobian matrix

$$J = \begin{bmatrix} \frac{\partial f}{\partial X} & \frac{\partial f}{\partial Y} \\ \frac{\partial g}{\partial X} & \frac{\partial g}{\partial Y} \end{bmatrix} = \begin{bmatrix} r_x \left(1 - 2X^* - a_{xy} Y^*\right) & -r_x a_{xy} Y^* \\ -r_y a_{yx} X^* & r_y \left(1 - 2Y^* - a_{yx} X^*\right) \end{bmatrix} \tag{25}$$

where $X' = f(X, Y)$ and $Y' = g(X, Y)$. For every, equilibrium point, we shall compute the Jacobian matrix at the specific equilibrium point and determine the trace ($\mathrm{Tr}(J)$), determinant ($\det(J)$) and discriminant $\Delta = \mathrm{Tr}^2(J) - 4\det(J)$ to provide a stabibility description for each point[2]. For the differential system (eqs. 5 and 6), there are four equilibrium points

[2]Please refer to the Poncaré diagram in the appendix to see the different classes of equilibrium points depending on the Jacobian matrix.

1. $\underline{(X_1^*, Y_1^*) = (0,0)}$ : we have

$$J_1 = \begin{bmatrix} r_x & 0 \\ 0 & r_y \end{bmatrix} \tag{26}$$

So $\mathrm{Tr}(J_1) = r_x + r_y > 0$, $\det(J_1) = r_x r_y > 0$ and $\Delta = r_x^2 r_y^2 > 0$, this means that this equilibrium point is unstable (source) and as a result, populations $X$ and $Y$ move away from it. Both populations cannot simultaneously be driven to extinction.

2. $\underline{(X_2^*, Y_2^*) = (1,0)}$ : we have

$$J_2 = \begin{bmatrix} -r_x & 0 \\ -r_y a_{yx} & r_y\,(1 - a_{yx}) \end{bmatrix} \tag{27}$$

So here we have

$$\begin{aligned} \mathrm{Tr}(J_2) &= -r_x + r_y\,(1 - a_{xy}) && (28) \\ \det(J_2) &= -r_x r_y\,(1 - a_{xy}) && (29) \\ \Delta &= [r_x + r_y(1 - a_{yx})]^2 && (30) \end{aligned}$$

We see that $\Delta > 0$, but there are two possible scenarios

- if $a_{yx} < 1$, then $\det(J_2) < 0$ and this point is an unstable saddle.
- if $a_{xy} > 1$, then $\det(J_2) > 0$ and $\mathrm{Tr}(J_2) < 0$, meaning this point is stable (sink). In this case, population $Y$ has been driven to extinction and $X$, having thrived, reaches its carrying capacity.

3. $\underline{(X_3^*, Y_3^*) = (0,1)}$ : this case is analog to the previous and the following conclusions can be drawn in a similar fashion

- if $a_{xy} < 1$, then $\det(J_3) < 0$ and this point is an unstable saddle.
- if $a_{yx} > 1$, then $\det(J_3) > 0$ and $\mathrm{Tr}(J_3) < 0$, meaning this point is stable (sink). In this case, population $X$ has been driven to extinction and $Y$, having thrived, reaches its carrying capacity.

4. $\underline{(X_4^*, Y_4^*) = \left(\frac{1-a_{xy}}{1-a_{xy}a_{yx}}, \frac{1-a_{yx}}{1-a_{xy}a_{yx}}\right)}$ : we have

$$J_4 = \begin{bmatrix} -r_x X^* & -r_x a_{xy} Y^* \\ -r_y a_{yx} X^* & -r_y Y^* \end{bmatrix} \tag{31}$$

So we find,

$$\begin{aligned} \mathrm{Tr}(J_2) &= -r_x X^* - r_y Y^* < 0 && (32) \\ \det(J_2) &= r_x r_y X^* Y^*\,(1 - a_{xy}a_{yx}) && (33) \\ \Delta &= r_x^2 X^{*2} + r_y^2 Y^{*2} - 2 r_x r_y X^* Y^*\,(2 - a_{xy}a_{yx}) && (34) \end{aligned}$$

Of course, since $(X_4^*, Y_4^*) \in \mathbb{R}_+^2$, either both $(a_{xy}, a_{yx})$ are simultaneously strictly larger than 1, or else simultaneously strictly smaller than 1

- if $a_{xy} < 1$ and $a_{yx} < 1$, then $\det(J_4) > 0$ and $\Delta > (r_x X^* - r_y Y^*)^2 > 0$. Therefore $(X_4^*, Y_4^*)$ is stable (sink) and this point is an equilibrium where both species coexist.
- if $a_{xy} > 1$ and $a_{yx} > 1$, then $\det(J_4) < 0$. As a result $(X_4^*, Y_4^*)$ is an unstable saddle point.

#### 4.3 Classical Predator-Prey model

In the classical predator-prey model, there are two equilibrium points

1. $\underline{(x_0^*, y_0^*) = (0,0)}$ : The Jacobian at this point is

$$J_0 = \begin{bmatrix} a & 0 \\ 0 & -d \end{bmatrix} \tag{35}$$

So $\det(J_0) < 0$ and this point is a saddle that cannot be reached from $\mathbb{R}_+^{*2}$. It can be approached on the $y$ axis (when $x = 0$).

2. $\underline{(x_1^*, y_1^*) = (\frac{d}{c}, \frac{a}{b})}$ : here the Jacobian is

$$J_1 = \begin{bmatrix} 0 & -\frac{bd}{c} \\ \frac{ac}{b} & 0 \end{bmatrix} \tag{36}$$

Since $\mathrm{Tr}(J_1) = 0$ and $\det(J_1) > 0$, $(x, y)$ solutions form closed orbits around this equilibrium point. The amplitude of the orbit depends on the initial conditions. A stable orbit necessarily exists as long as the initial conditions $x_0$ and $y_0$ are simultaneously non-zero.

#### 4.4 Predator-Prey model with carrying capacity

In this case, there are three equilibrium points.

1. $\underline{(x_0^*, y_0^*) = (0,0)}$ : here, the Jacobian is identical to that of the classical predator-prey model and this point is a saddle that can only be approached from the y-axis if $x = 0$.

2. $\underline{(x_1^*, y_1^*) = (\kappa, 0)}$ : here the Jacobian is

$$J_1 = \begin{bmatrix} -a & -b\kappa \\ 0 & c\kappa - d \end{bmatrix} \tag{37}$$

If $\frac{d}{c\kappa} < 1$, then $\det J_1$ is a saddle point that can only be reached from the x-axis when $y = 0$. If $\frac{d}{c\kappa} \geq 1$,then this point is a stable sink. This is a case of course, where the predators would go extinct and the prey reach their carrying capacity. In this case, similarly to the single population competitive case, the final death rate is $m_1' = a\kappa$

3. $\underline{(x_2^*, y_2^*) = (\frac{d}{c}, \frac{a}{b}[1 - \frac{d}{\kappa c}])}$: of course, since there are no negative populations, this point can only exist under the condition that $\frac{d}{c\kappa} < 1$, which is the saddle point condition of $(x_1^*, y_1^*)$. The Jacobian reads

$$J_2 = \begin{bmatrix} -a\frac{d}{c\kappa} & -\frac{bd}{c} \\ \frac{ac}{b}(1 - \frac{d}{c\kappa}) & 0 \end{bmatrix} \tag{38}$$

Since $\det(J_2) > 0$ and $\mathrm{Tr}(J_2) < 0$, this point is a sink and corresponds to a stable coexistence scenario. Here the final death rate is

$$m_2' = ad\left(\frac{1}{b} + \frac{1}{c} - \frac{d}{cb\kappa}\right) \tag{39}$$

### References

[1] Fred Brauer, Carlos Castillo-Chavez, and Carlos Castillo-Chavez. *Mathematical models in population biology and epidemiology*, volume 2. Springer, 2012.
